# Correspondence between sleep patterns and anhedonia in adult male mice exposed to early-life stress

**DOI:** 10.1101/2025.04.18.649529

**Authors:** Andrew Hall, Dominika J. Burek, Erin E. Hisey, Lauren T. Seabrook, Emma L. Fritsch, Zoe G. Beatty, Kerry J. Ressler, Brian D. Kangas, William A. Carlezon

## Abstract

Early-life stress (ELS) can produce long-lasting effects that increase the risk for mood and anxiety disorders. Transdiagnostic symptoms include anhedonia (reduced reward sensitivity) and sleep disruption, both of which are quantifiable via objective endpoints that can be utilized across species. Here we used a mouse model for ELS—exposure to juvenile chronic social defeat stress (jCSDS)—together with translationally-applicable endpoints to examine correspondence between sleep patterns and anhedonia. These initial studies focused on males, which typically show robust defeat-induced anhedonia phenotypes. Exposure to jCSDS produced reductions in open-field social behavior, an endpoint commonly used in mice to quantify stress effects, during adulthood. Mice were then implanted with wireless transmitters that enable continuous EEG-derived analysis of sleep architecture. Following assessment of baseline sleep patterns, mice were tested in a rodent version of the Probabilistic Reward Task (PRT), a procedure used to quantify reward responsiveness in humans, during the light phase of their diurnal cycle. These studies revealed significant associations between baseline sleep architecture and anhedonic phenotypes in jCSDS-exposed mice: higher anhedonia correlated with less time awake and more time in slow wave sleep (SWS) during the light phase, and more time awake and less time in rapid eye movement (REM) sleep and SWS during the dark (active) phase. Our findings suggest that sleep patterns represent a biomarker that can predict stress-susceptible (higher anhedonia) and resilient (lower anhedonia) phenotypes. This work enhances our understanding of relationships between sleep and anhedonia, and may provide a basis for precision approaches to treat ELS-induced pathophysiology.

## INTRODUCTION

Stress can cause or exacerbate a wide range of psychiatric illnesses [*1–6*]. The signs and symptoms of these illnesses often overlap, with numerous conditions sharing core features including anhedonia (reduced sensitivity to reward) and sleep dysregulation (sleeping too much, too little, or altered patterns of sleep) [*1,7*]. Indeed, stress is known to disrupt sleep, which exacerbates the signs of psychiatric illness [*1,8–14*]. Importantly, some individuals are especially sensitive to stress while others appear resilient, which has created interest in identification of stress biomarkers (objective indicators of a biological condition). The ability to use biomarkers to more reliably identify individuals that have, or are likely to develop, symptoms of stress-related psychiatric illness will enable precision psychiatry [*15*] and improved mental health outcomes.

In humans, it is established that early-life stress (ELS)—encompassing conditions including child abuse, bullying, neglect, and poverty/low resource environments—is a risk factor for developing mood and anxiety disorders [*16,17*]. ELS affects approximately 1 in 7 children and is associated with alterations in brain connectivity and anatomy during adolescence and adulthood [*16–18*]. While rodent models of early-life neglect and/or caregiver maltreatment [*19–24*] have been developed, there are few reports describing the effects of ELS with a physical component [*25,26*]. In mice, adult chronic social defeat stress (aCSDS)—which includes both physical (defeat) and emotional (sharing a cage with an aggressive/dominant conspecific) elements—is a commonly used model of stress that produces alterations in endpoints including social behavior, motivation, and sleep [*13,14,26–31*]. Complex behavioral tests such as intracranial self-stimulation (ICSS) unequivocally demonstrate that aCSDS produces anhedonia [*30–32*], reflected by reductions in the rewarding effects of lateral hypothalamic brain stimulation without changes in response capacity, but have no complementary paradigm in humans to validate translational relevance. Approaches that are translationally-relevant—enabling use of the same procedures and endpoints across species—are needed to improve the ability of models to predict outcomes in humans. We developed a juvenile chronic social defeat stress (jCSDS) procedure that produces rigorous and persistent changes in behavior during adulthood as the result of early-life stress [*33,34*], and paired it with a touchscreen-based mouse version of the Probabilistic Reward Task (PRT), a translationally-relevant behavioral procedure that is used in humans to study anhedonia and/or to evaluate the efficacy of therapeutics [*35–37*]. Using the PRT, we found that jCSDS-exposed mice showed behavioral patterns that are considered to reflect anhedonia when seen in humans [*33,35–37*]. The ability to align test procedures and endpoints across species is an important step toward developing biomarkers in mice that are translationally-relevant and scalable, while having maximal predictive value in humans.

Sleep has advantages as a translationally-relevant biomarker because it is measured and defined in the same way across species [*2,38*]. Stress-induced changes in sleep patterns and microstructure (architecture) have been reported in both humans and mice [*13,14,39–44*], and are considered a maladaptive outcome of ELS in humans [*8*]. Sleep-related endpoints are complex, and there can be major differences in sleep pattens even within a diagnostic classification: indeed, depressive illness can be associated with either increases or decreases in sleep [*1*], suggesting that dysregulation of normal sleep patterns—regardless of direction—is currently the defining feature. While certain sleep characteristics are associated with risk for the development and/or relapse of depression [*45–47*], little is known about specific relationships between sleep patterns and any individual depression symptom, including motivational deficits (anhedonia). Importantly, anhedonia is not a diagnosis but rather a symptom that cuts across numerous psychiatric conditions [*1,7*]; while it is often considered to be one of the most debilitating symptoms of depression, it does not define the condition and there are experimental approaches that treat it specifically [*35,37*]. Considering the accessibility of devices (e.g., smartphones and watches) that provide sleep metrics, a better understanding of relationships between sleep biomarkers and motivation may enable rapid improvements in the diagnosis, treatment, and prevention of myriad forms of psychiatric illness [*14,48*].

The present studies were designed to determine if ELS in mice causes changes in sleep patterns during adulthood that correspond with changes in motivated behavior. Initial studies focused on males because the effects of CSDS on reward function have been challenging to reliably document in female rodents [*49*]. While key elements were based upon our prior work, the overall experimental design was novel. Our data indicate that this design produced some findings that differ from those in previous reports—namely, differences between the effects of aCSDS and jCSDS on sleep [*13*], and increased variability in PRT performance in mice implanted with wireless transmitters [*33*]—but also revealed strong associations between baseline sleep patterns and motivated behavior.

## METHODS

### Mice

Subjects were C57BL/6J (C57) mice obtained at 3 weeks of age from Jackson Laboratory (Bar Harbor, ME, USA). To generate aggressors, virgin male and ovariectomized female Swiss Webster (CFW) mice were obtained at 8 weeks of age from Charles River (Shrewsbury, MA, USA). All mice were housed under a 12-h light and 12-h dark schedule (07:00-on, 19:00-off) in a temperature- (21 ± 2.0 °C) and humidity-controlled environment (50 ± 20%), and food and water were available *ad libitum* until PRT training. Subject mice were given one week to habituate prior to study onset. All experimental procedures were approved by the Institutional Animal Care and Use Committee at McLean Hospital and were performed in accordance with the National Institutes of Health’s (NIH) Guide for the Care and Use of Animals.

### Juvenile chronic social defeat stress (jCSDS)

Virgin male CFW mice were housed with ovariectomized female CFW mice for at least one week, then screened for aggression in interactions with non-experimental C57 male mice. Only male CFWs that showed an attack latency of <30 seconds for 2 consecutive days were used for defeat sessions. Female CFW mice were permanently removed from the cage after screening. Beginning on postnatal day (P)30 (± 2 days), each C57 mice was subjected to a 10-day jCSDS regimen, as described previously [*33,34*]. To begin each daily session, a C57 (subject) mouse was placed into the home cage of a male CFW (aggressor) mouse. Each session proceeded until the aggressor had delivered 30 bites or 5 min had elapsed. After each defeat session, the mice were separated in the cage with a perforated Plexiglas divider, allowing continuous visual and olfactory contact without further physical interactions. This procedure was repeated for 10 consecutive days, with each day involving a new cage and aggressor. Control mice were handled daily and housed in identical cages as the defeated mice but instead opposite a conspecific (C57) mouse [*29–31*].

### Open field social interaction (OFSI)

To examine the persistence of ELS effects on behavior, mice that received jCSDS were tested for OFSI during adulthood, as described [*33*]. On P70 (± 2 days), mice were transported from the vivarium to a behavioral testing room, where they habituated for one hour. Each C57 mouse was then placed in a square arena (45.7 x 45.7 cm) containing an empty wire cup for 150 sec. The C57 was then removed, and a non-aggressive CFW male was placed under the wire cup. The C57 was then placed back in the arena for an additional 150 sec. Social interaction ratio was calculated as the amount of time spent in the social interaction zone (2-cm circular zone around the wire cup) when the CFW was inside the cup divided by the amount of time spent in this area when the cup was empty.

### EEG- and EMG-based sleep telemetry: transmitter implantation surgeries

To examine the persistence of ELS effects on sleep, mice were implanted with wireless telemetry devices (HD-X02; DSI, St. Paul, MN, USA) to enable continuous recording of electroencephalograms (EEG), electromyograms (EMG), locomotor activity, and subcutaneous (SC) temperature. Surgery was performed on P72 (± 2 days) using procedures described previously [*13*]. Briefly, mice were anesthetized with 2% isoflurane and immobilized on a stereotaxic instrument. A small incision was made from the medial skull to posterior neck, through which a subcutaneous pocket on the back was opened using lubricated forceps. Transmitters were inserted lateral to the spine, midway between the fore and hind limbs. Two incisions were made in the trapezius muscle using a 21-G needle, through which EMG electrodes were threaded and secured with sutures. EEG electrodes were attached to two stainless steel screws, inserted into the skull, and lowered until they contacted dura (relative to bregma: frontal AP=+1.0 mm, ML=+1.0 mm; parietal AP=−3.0 mm, ML=−3.0 mm). Screws and EEG leads were secured with dental cement and the incision was closed with sutures. Triple antibiotic ointment (topical) and lidocaine (2%, topical) were applied to the incision site. Ketoprofen (5.0 mg/kg, SC) was administered immediately after surgery, and once daily for 7 days of recovery and post-operative monitoring with food and hydration supplements.

### Sleep telemetry: data acquisition and analyses

Mice were single housed in N10 mouse cages on a standard 12h-light/12h-dark cycle. Cages sat on RPC-1 PhysioTel receiver platforms (DSI), which detect signals emitted by the HD-X02 transmitters. Receivers connected to a data exchange matrix that continuously uploaded recording data (sampling rate=500 Hz) to Ponemah Software (DSI), which were analyzed in Neuroscore Software V3.4 (DSI). Raw EEG signal was quantified into relative spectral power in 10-sec epochs with Fast Fourier Transform (FFT) and a Hamming signal window. Frequency bands were defined as delta (0.5–4 Hz), theta (4–8 Hz), alpha (8–12 Hz), beta (16–24 Hz), and low gamma (30–50 Hz). Vigilance states of active wakefulness (AW), REM sleep, and SWS were manually assigned. For any 10-sec epoch, AW required high EMG activity; REM required largest peak in theta power with negligible EMG activity; and SWS required largest peak in delta power with negligible EMG activity. Total duration and uninterrupted bouts of each stage were calculated separately for 12 hr lights-on/12 hr lights-off per 24-hr period. Baseline was averaged from 3 consecutive days of continuous recording (P85-87, ± 2 days) immediately prior to starting the PRT studies.

### Probabilistic Reward Task (PRT): training and testing

The PRT is a touchscreen task reverse-engineered from human studies and used in mice as described previously [*33,36*]. On P88 (± 2 days)—3 days before PRT training—mice were food-restricted to 85% of their free-fed body weight. Mice were first trained to rear and touch a 5 x 5 cm blue square on a black background, in various positions on the touchscreen, to receive 0.02 mL of a highly palatable 20% sweetened condensed milk reward, which was delivered in a well on the opposite wall of the touchscreen. Mice were then trained during 100-trial sessions to discriminate between a long or short white line (24 x 3 cm or 12 x 3 cm) on a black background by responding on one of two virtual levers (5 x 5 cm blue squares) presented below the line, to the left and right of center. Correct responses were rewarded with sweetened condensed milk paired with a tone and brief, bright yellow screen followed by a 10-sec blackout period, whereas incorrect responses resulted in a 20-sec timeout. During initial PRT training sessions, a correction procedure was employed, where incorrect trials were repeated until a correct response was made prior to advancing to the next trial. After reaching criterion in this phase (≤10 errors for both the long and short lines on consecutive sessions), mice were tested without correction under otherwise identical contingencies. Upon reaching >80% accuracy during two consecutive sessions without correction, which generally required 3-4 weeks, PRT testing commenced. Mice were exposed to a 5-session PRT testing protocol using 3:1 probabilistic reinforcement contingencies, such that a correct response to one of the line lengths (long or short) was reinforced 60% of the time (rich stimulus), whereas a correct response to the other line length was reinforced 20% of the time (lean stimulus). Incorrect responses were never reinforced. The line length associated with the rich and lean contingency was determined for each subject during their final two line-length discrimination training sessions, by examining their accuracies and designating the line length with a higher mean accuracy as the stimulus to be rewarded on the lean schedule. This approach was expressly designed to examine the effects of jCSDS on response bias generated by responsivity to asymmetrical probabilistic contingencies, rather than the amplification of a preexisting inherent bias that is a function of uncontrolled variables [*33,36*].

### Statistical analyses

All analyses were performed using GraphPad Prism 9 software (San Diego, CA), with significance set to *P*<0.05. For OFSI, mean social interaction ratios were assessed for normality with Kolmogorov-Smirnov tests, and then compared with unpaired t-tests. Likewise, for sleep studies, sleep stage durations and bouts over the 3-day analyses period were assessed for normality with Kolmogorov-Smirnov tests and then compared with unpaired t-tests. Home cage locomotor activity was quantified using continuous data collection over the same period and analyzed similarly. For the PRT, test sessions yield two primary dependent measures: response bias (log *b*, which reflects reward responsiveness) and task discriminability (log *d,* which reflects discriminative abilities in the task). These values are quantified using equations derived from signal detection theory by examining the number of correct and incorrect responses for rich and lean trial types; detailed descriptions of the calculations and their derivation have been described previously [*33,36*]. High bias values are produced by high numbers of correct responses for rich trials and incorrect responses for lean trials, whereas high discriminability values are produced by high numbers of correct responses for both rich and lean trials. Relationships between sleep stage duration and log *b* were determined with simple linear regressions, calculating 95% confidence bands of the best-fit line and comparing whether slope and intercepts differed from 0 (within-group) or between control and jCSDS-exposed mice (between-group). Group differences in slope were assessed using analyses of covariance (ANCOVAs).

## RESULTS

The experimental flow is depicted in **Fig. 1**. At the outset of the experiment, there were 7 jCSDS-exposed mice and 7 controls. Following recovery from sleep transmitter implantation surgery, one of the jCSDS-exposed mice showed overt signs of infection including lethargy that were accompanied by abnormally high body temperature readings. This mouse was excluded from all analyses.

**Fig. 1.**
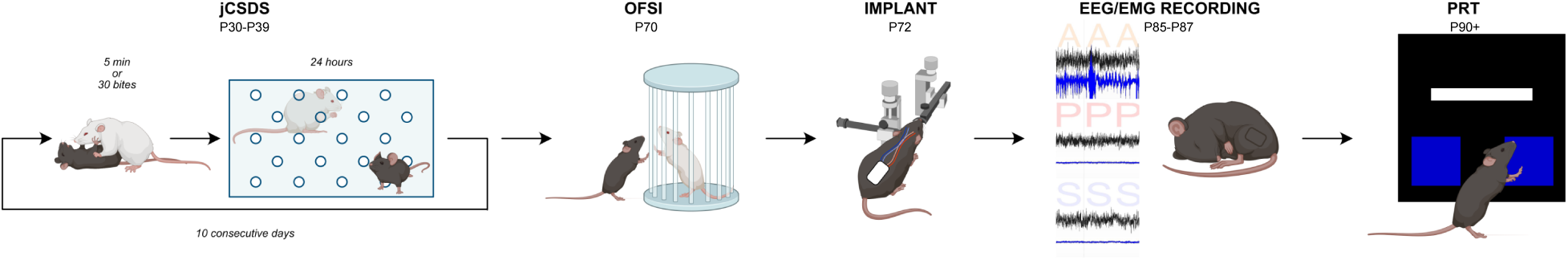
Schematic of the experimental design. Juvenile (j)C57BL/6J male mice were defeated by adult CFW male aggressors each day on 10 consecutive days. Each defeat session was followed by 24 hr of co-housing separated by a perforated plastic divider. To determine the persistence of jCSDS-induced behavioral effects, the mice were assessed during adulthood alterations in social behavior (in open field social interaction tests [*OFSI*]), sleep architecture (via wireless EEG/EMG telemetry), and reward sensitivity (in the Probabilistic Reward Task [*PRT*]).

### OFSI and baseline home-cage locomotor activity

We first examined the effects of jCSDS on social interaction in OFSI tests performed during adulthood (on P70). As reported previously [*33*], the social interaction ratios of jCSDS-exposed mice were significantly lower than those of controls (t_11_=5.779, p=0.0001) (**Fig. 2A**). Likewise, jCSDS-exposed mice had significantly lower raw social interaction time compared to controls (t_11_=2.855, p=0.0157) (**Fig. 2B**). Confirming previous findings, there were no group differences in overall locomotor activity during the OFSI sessions (not shown), and a more comprehensive analysis involving continuous collection of baseline home-cage locomotor activity data via wireless telemetry over an entire 3-day period (P85-87) also indicated no group differences in average daily locomotor activity counts (t_11_=0.4712, not significant [*n.s.*]) (**Fig. 2C**). These findings suggest that jCSDS-exposed mice retain a long-lasting memory of ELS into adulthood, creating persistent signs of social avoidance behavior without more general deficits in activity.

**Fig. 2.**
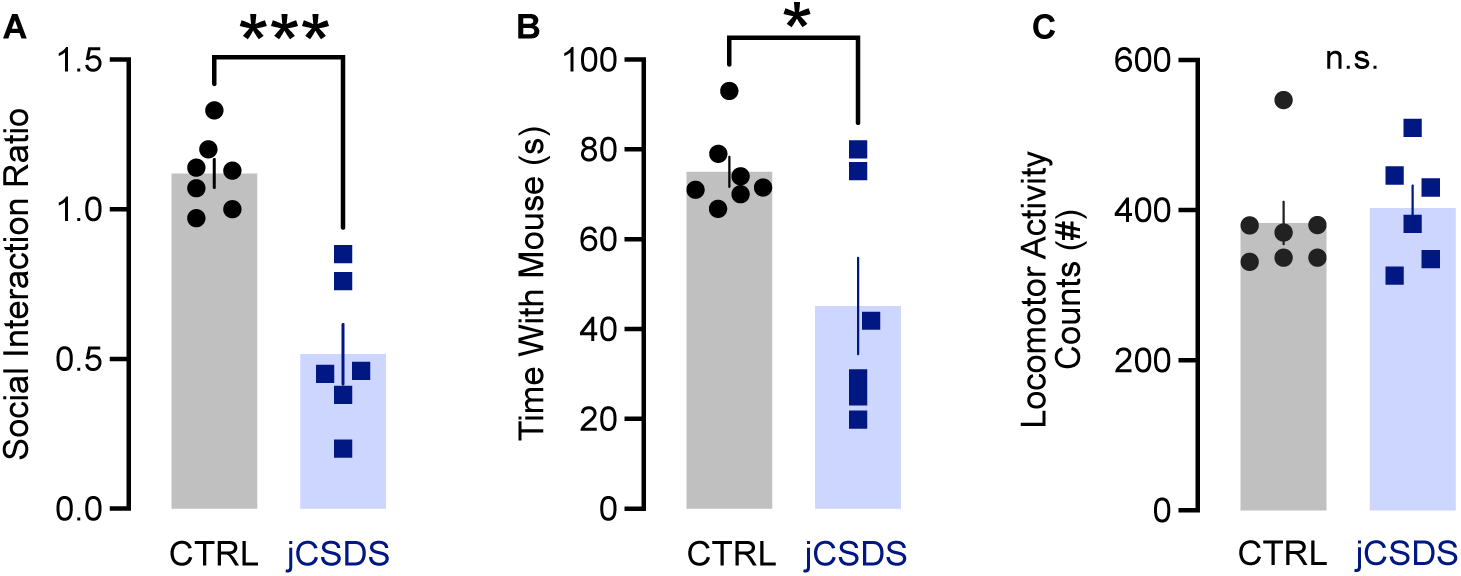
Behavior in open field social interaction tests (P70) and home cage locomotor activity (P85-87). (**A**) Ratio of interaction duration with empty cup versus with novel CFW mouse. (**B**) Interaction duration with novel CFW mouse. (**C**) Mean daily locomotor activity counts in the home cage over a 3-day period (P85-87). \**P*<0.05, \*\*\**P*<0.001, n.s. (not significant), Student’s t-tests.

### Baseline sleep

Following >10 days recovery from transmitter implantation surgery, we examined the effects of jCSDS on sleep patterns, using continuous EEG/EMG data to compare average time spent awake, in REM sleep, and in SWS. Each 24-hr day was split into a 12-hr lights-on period (when mice are more likely to be asleep) and 12-hr lights-off period (when mice are more likely to be active). There were no group differences in any of these endpoints during either the light or dark phase. During the light phase, there were no differences between jCSDS-exposed mice and controls in wake duration (t_11_=0.9971, n.s.) (**Fig. 3A**), REM duration (t_11_=0.3590, n.s.) (**Fig. 3B**), or SWS duration (t_11_=0.7145, n.s.) (**Fig. 3C**). Likewise, during the dark phase there were no group differences in wake duration (t_11_=0.2677, n.s.) (**Fig. 3D**), REM duration (t_11_=0.8199, n.s.) (**Fig. 3E**), or SWS duration (t_11_=0.2778, n.s.) (**Fig. 3F**). There were no effects seen on bouts of these vigilance states during either phase of the light cycle (not shown). As secondary measures, there were also no group differences in mean body temperature or locomotor activity (all t-values <0.7, not shown). These data indicate that, at this level of analysis, ELS does not have broad effects on sleep patterns, body temperature, or locomotor activity that persist into adulthood.

**Fig. 3.**
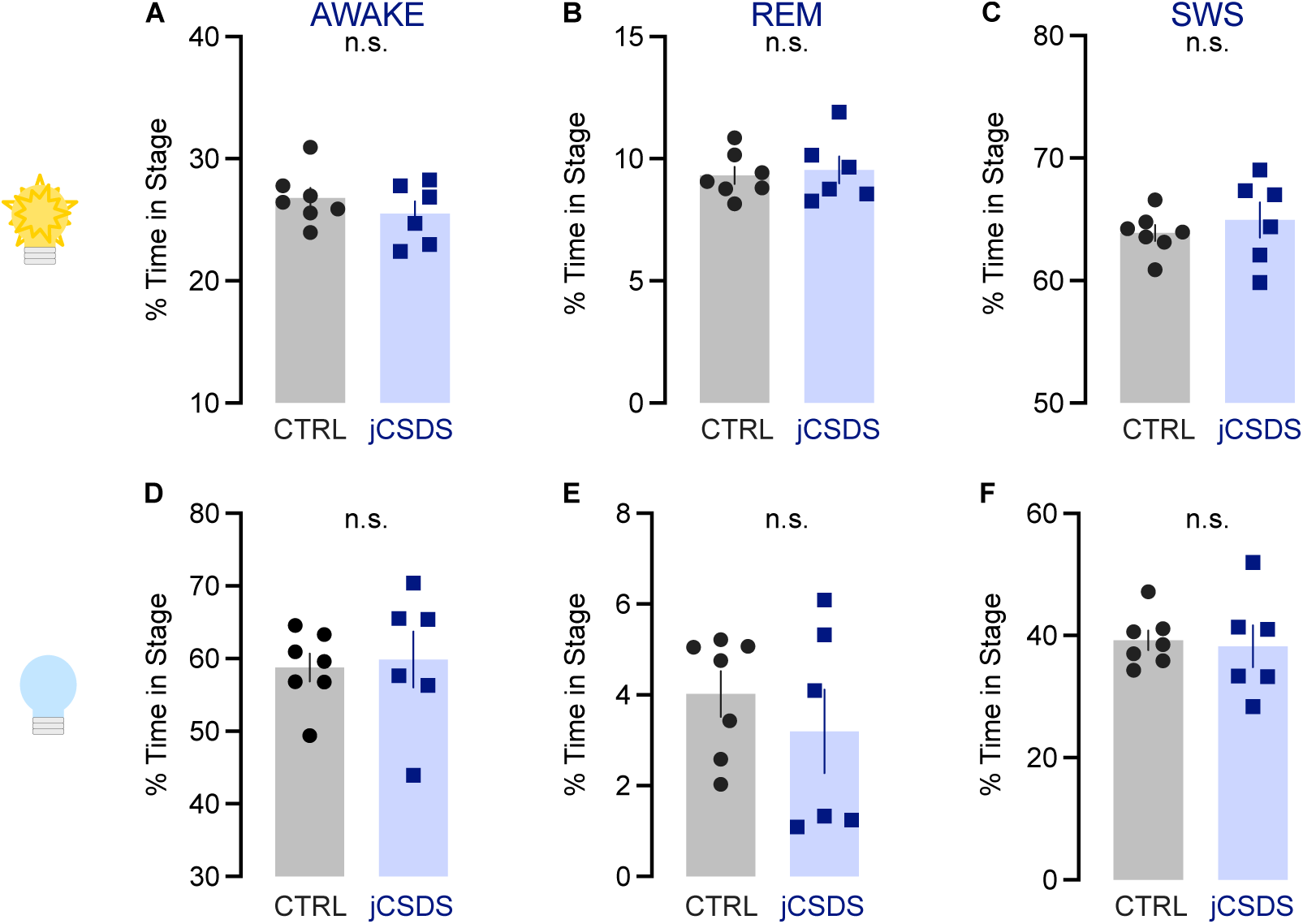
Assessment of mean baseline sleep architecture across days P85-87. During the 12-hr lights-on phase of the daily light cycle (top row), (**A**) time spent awake, (**B**) in REM sleep, and (**C**) in slow wave sleep (SWS). During the lights-off phase (bottom row), (**D**) time spent awake (**E**) in REM, and (**F**) in SWS. Not significant (n.s.), Student’s t-tests.

### PRT: reward responsiveness and discriminative ability

Mice were then trained and tested in the PRT for reward responsiveness (quantified by log *b*) and discriminative ability (quantified by log *d*). In contrast to a previous report where we found that jCSDS-exposed mice showed significant deficits in reward responsiveness at a similar timepoint [*33*], there were no group differences in log *b* (t_11_=0.5927, n.s.) (**Fig. 4A**). While some mice showed reductions in log *b*, the distribution of responses was larger than in previous studies. Consistent with our previous report, we also observed no significant difference in log *d* (t_11_=1.633, n.s.) (**Fig. 4B**). While the failure to replicate previous PRT findings was surprising, the differences may be attributable to the fact that mice received an intervening procedure between the OFSI and PRT (surgical implantation of wireless transmitters for sleep analyses) in the present studies.

**Fig. 4.**
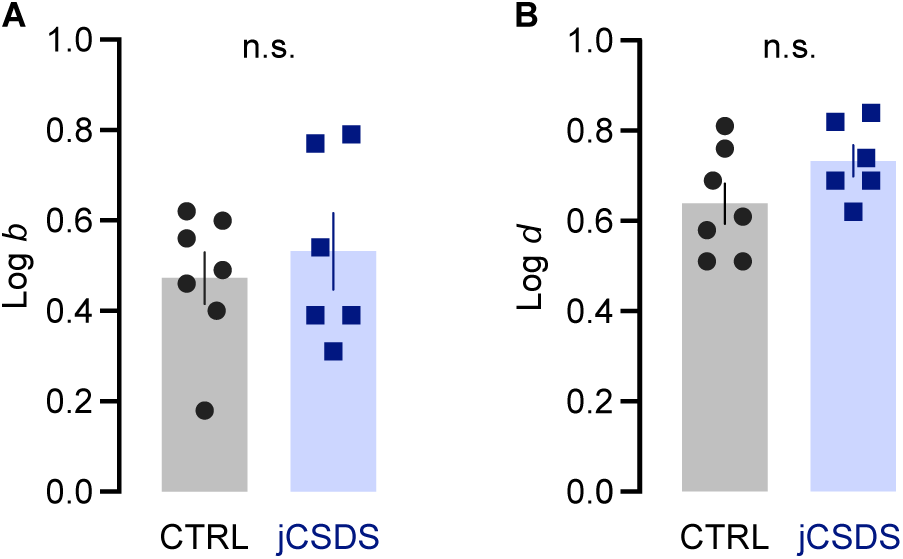
Mean performance in probabilistic reward task (PRT) across 5 consecutive days of testing. Training began on P90. (**A**) Response bias (log *b*), and (**B**) task discriminability (log *d*). Not significant (n.s.), Student’s t-tests.

### Correlations between sleep and reward function: light phase

The increased variability in log *b* in jCSDS-exposed mice offered an opportunity to examine correlations between sleep and PRT endpoints. Regression analyses revealed correlations between log *b* and vigilance states in jCSDS-exposed mice, whereas no associations were observed in controls (i.e., the slopes never differed from 0). During the light phase (when mice are less likely to be active), there was a significant positive correlation in jCSDS-exposed mice between time spent awake and log *b* (Y=11.11X+19.60, F_(1,4)_=21.51, p=0.0097), such that the mice with lower log *b* values tended to spend a lower percentage of time awake (i.e., the more anhedonic mice spent more time asleep). While this correlation was statistically significant, there were no group differences in slopes (F_(1,9)_=1.539, n.s.) (**Fig. 5A**). There were no correlations between time spent in REM and log *b* (**Fig. 5B**), but there was a significant negative correlation in jCSDS-exposed mice between time spent in SWS and log *b* (Y=-15.8X+73.36, F_(1,4)_=26.29, p=0.0068), such that the mice with lower log *b* values tended to spend a higher percentage of time in SWS (i.e., the more anhedonic mice spent more time in SWS), with significant differences in slopes (F_(1,9)_=5.597, p=0.0422) (**Fig. 5C**). There was also a negative correlation between SWS bouts and log *b* (Y=-2.673X+51.71, F_(1,4)_=9.708, p=0.0357), with no group differences in slopes (not shown); parallel changes in SWS time and bouts suggests minimal fragmentation. These findings suggest that more total time asleep—and in SWS specifically—during the light phase is a biomarker that can predict ELS-induced anhedonia.

**Fig. 5.**
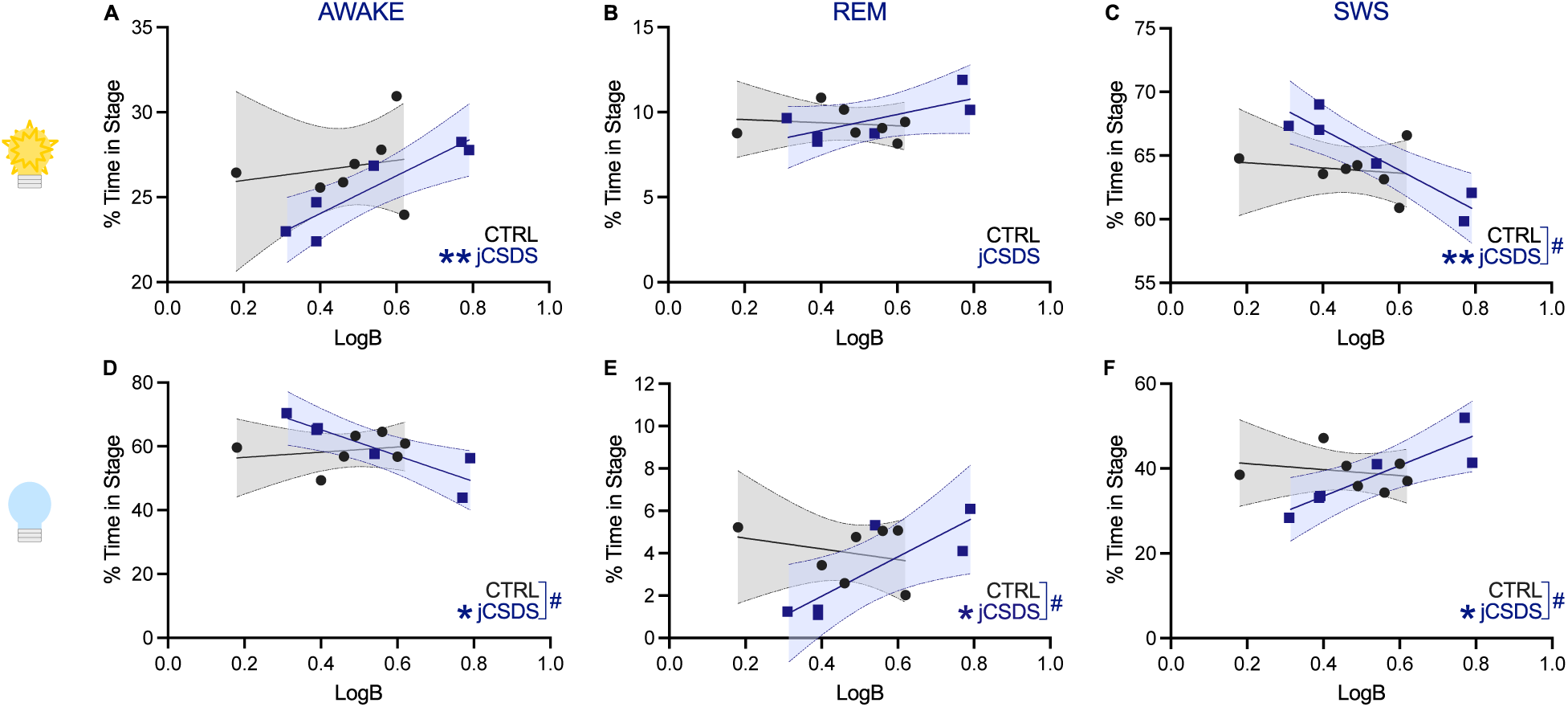
Regression analyses of group differences in correlations between mean baseline sleep data and mean log *b* in the Probabilistic Reward Task (PRT). Lower log *b* data indicate higher levels of anhedonia. During the 12-hr lights-on phase of the daily light cycle (top row), correlations between log *b* and time spent (**A**) awake, (**B**) in REM, and (**C**) in slow-wave sleep (SWS). During the 12-hr lights-off phase (bottom row), correlations between log *b* and time spent (**D**) awake, (**E**) in REM, and (**F**) in SWS. \**P*<0.01, \*\**P*<0.01, linear regression analyses; #*P*<0.05, analyses of covariance (ANCOVA).

### Correlations between sleep and reward function: dark phase

During the dark phase (when mice are more likely to be active), there was a significant negative correlation in jCSDS-exposed mice between time spent awake and log *b* (Y=- 40.66X+81.48, F_(1,4)_=15.10, p=0.0178), such that the mice with lower log *b* values tended to spend a higher percentage of time awake (i.e., the more anhedonic mice spent less time asleep), with statistically significant group differences in slopes (F_(1,9)_=7.430, p=0.0234) (**Fig. 5D**). There was a positive correlation with time spent in REM (Y=9.287X- 1.743, F_(1,4)_=10.36, p=0.0323), such that the mice with lower log *b* values tended to spend a lower percentage of time in REM (i.e., the more anhedonic mice spent less time in REM), with statistically significant group differences in slopes (F_(1,9)_=6.2549, p=0.0338) (**Fig. 5E**). There was also a positive correlation between REM bouts and log *b* (F_(1,4)_=8.271, p=0.0452), with no group differences in slopes (not shown); parallel changes in REM time and bouts suggests minimal fragmentation. In addition, there was a significant positive correlation in jCSDS-exposed mice between time spent in SWS and log *b* (Y=36.09X+19.05, F_(1,4)_=14.85, p=0.0183), such that the mice with lower log *b* values tended to spend a lower percentage of time in SWS (i.e., the more anhedonic mice spent less time in SWS), with significant differences in slopes (F_(1,9)_=7.861, p=0.0206) (**Fig. 5F**). These findings suggest that more total time awake, with correspondingly less time in REM and SWS, during the dark phase is a biomarker that can predict ELS-induced anhedonia.

## DISCUSSION

We report that ELS in male mice produces associations between sleep patterns and anhedonia, a common feature of depressive illness, during adulthood. To produce ELS, we used jCSDS, which we have previously shown causes social avoidance behavior as well as signs of anhedonia in a rodent version of the PRT [*33,36*], a touchscreen-based procedure used in humans to assess reward responsiveness [*35,37*]. Consistent with our previous report [*33*], mice exposed to jCSDS from PD30-39 showed strong and persistent social avoidance phenotypes in OFSI tests conducted on P70, which is a characteristic outcome of CSDS [*26–30*] that indicates efficacy of the stress protocol. When sleep architecture was later assessed in jCSDS-exposed mice over a 3-day baseline period during adulthood (∼P86) via a wireless EEG/EMG system, there were no major differences in wakefulness, REM, or SWS when compared to mice exposed to control conditions. Likewise, when jCSDS-exposed mice were subsequently tested in the PRT, there were no differences in log *b* (a measure of reward responsiveness) or log *d* (a measure of response capabilities) [*33,36*]. The lack of an overall effect of in the PRT is inconsistent with our previous report that jCSDS causes behavioral patterns in this procedure during adulthood that are considered to reflect anhedonia when seen in humans [*33,35–37*]. However, whereas the jCSDS-exposed mice in previous studies showed relatively uniform behavioral responses in the PRT, the mice in the present studies—which had undergone an intervening surgical procedure to enable collection of EEG/EMG-based sleep data—showed more individual variability, covering a range of high and low levels of anhedonia. This variability enabled use of regression analyses to reveal novel relationships between baseline sleep patterns and anhedonia. Specifically, higher anhedonia was correlated with less time awake and more time in SWS during the light (less active) phase, and more time awake and less time in REM and SWS during the dark (more active) phase. There were no correlations between these vigilance states and anhedonia in control mice, regardless of light-cycle phase. These findings suggest that ELS triggers adaptations that strengthen the link between sleep patterns and motivated behavior, and more broadly, that sleep patterns represent a biomarker that can differentiate stress-susceptible (higher anhedonia) and resilient (lower anhedonia) phenotypes.

These findings are consistent with numerous previous reports. We previously examined the effects of aCSDS on sleep architecture in real time, and found acute effects during the 10-day defeat regimen as well as longer-lasting effects that persisted throughout a 5-day recovery period [*13*]. Using analyses that covered the entire 24-hr daily light cycle, the most prominent effects of aCSDS were decreases in wakefulness accompanied by increases in the time spent in SWS and REM as well as increases in the number of REM bouts. These effects on sleep developed and peaked over a time course that is virtually identical to that seen in a parallel ICSS study in which anhedonia was quantified [*30*], suggesting links between these sleep patterns and anhedonia. Although the effects on wakefulness and time in REM quickly recovered, the increases in SWS time and REM bouts persisted throughout the recovery period, suggesting long-term effects on these endpoints. Effects seen in the present studies suggest that jCSDS produces long-lasting increases in SWS—evident ∼7 weeks after the jCSDS regimen and accompanied by decreases in wakefulness—in mice that subsequently show the highest levels of anhedonia (i.e., lowest log *b*), although inclusion of light-cycle phase isolates this effect to the light phase, when mice are normally less likely to be active. Interestingly, similar light cycle-dependent effects were seen during adulthood in mice exposed to another form of ELS: perinatal immune activation (PIA). Specifically, mice exposed to a PIA regimen that causes depressive- and anxiety-like behaviors [*50*] also show decreases in wakefulness and increases in SWS time during the light phase, with no changes during the dark phase, that are stable and persistent over the lifespan [*51*]. These findings are broadly consistent with findings of increased SWS in humans with depressive illness, although this is sometimes considered an adaptive response that serves to mitigate (rather than exacerbate) the debilitating symptoms of the condition; indeed, many standard antidepressant drugs increase SWS [*11*]. Interestingly, separating the sleep data into light-cycle phases revealed a complementary pattern of effects during the dark phase, when mice are normally more likely to be awake: mice with higher anhedonia spent more time awake and correspondingly less time in SWS and REM. While cause-effect relationships are difficult to establish in the experimental design we used for the present studies, one possibility is that these increases in wakefulness/reduced sleep during more active phases contribute to corresponding decreases in wakefulness and increased sleep during less active phases, resulting in persistent dysregulation of sleep architecture. In humans, insomnia is associated with mood and anxiety disorders [*11*], although this construct is not easily applied to mice because their sleep is distributed throughout the day, with periods of being more or less likely to be active as opposed to periods where sleep and wakefulness typically occur.

Some findings from the present studies were surprising based on the published literature, although the discrepancies are likely related to differences in experimental design. As one example, we previously reported that jCSDS produces increases in anhedonia (reflected by reductions in log *b*) during adulthood [*33*], whereas in the present studies the effect was not significant due to higher variability. However, there are important differences in the present experimental design and the previous work: foremost, the mice in the present study underwent a surgical procedure to implant the wireless transmitters before PRT training and testing. This finding raises the possibility that this additional experience increases variability and exacerbates individual differences and separation into susceptible and resilient populations. The addition of the sleep studies also resulted in the PRT studies starting ∼2 weeks later than in our original report. Importantly, previous work showed that there is no correlation between social behavior in the OFSI and reward-related behavior in the PRT, suggesting that these tests assess different domains [*7,33*]. In addition, previous studies of aCSDS showed acute effects on time spent in REM sleep and REM bouts [*13*] during periods where parallel studies showed strong anhedonia in the ICSS test [*30*], with the effect of bouts persisting through the brief (5-day) recovery period. In the present studies we did not observe correlations between increases in these REM-related endpoints and elevated anhedonia, but there are two major differences in the experimental design: the mice were defeated as juveniles rather than adults, and sleep was assessed weeks after the defeat regimen rather than during and immediately afterwards. These findings raise the possibility that REM effects are less persistent following ELS, which can be addressed in future studies. While increases in REM sleep in humans have been associated with elevated risk for relapse to depression [*52*], it is important to note that this risk has not been specifically tied to anhedonia or any other symptom. As described above, anhedonia is not a diagnosis but rather a symptom that cuts across many forms of psychiatric illness, and it can be more prevalent in some individuals than others. For example, anhedonia is particularly prevalent in the melancholic subtype of depression [*53,54*]. Our data raise the possibility of specific relationships between sleep and anhedonia, providing testable hypotheses that can be explored in humans experiencing anhedonia as a predominant feature of illness.

These early, proof-of-principle studies provide a basis for future work across species that focuses specifically on the relationship between sleep patterns and anhedonia. In humans, such studies might focus on determining whether anhedonia—as opposed to a broader diagnosis of depression—correlates with any of the sleep patterns identified in the present studies. The prediction would be that anhedonia would be highest in individuals with imbalance in sleep patterns characterized by both (i) more time asleep and in SWS during periods when sleep normally predominates (night) and (ii) less time asleep and less time in REM and SWS during periods when sleep can occur but is normally less prevalent (day). These types of studies may be facilitated by enriching subject selection with individuals with the melancholic subtype of depression [*54*] and using digital devices to track sleep on a continuous basis [*48*]. Studies in animal model systems could examine overlap in circuits that regulate sleep and motivated behavior. Indeed, we have previously reported that chemogenetic manipulations in the nucleus accumbens (NAc), a brain area implicated in motivated behavior, can affect sleep patterns and create phenotypes resembling those seen after aCSDS [*14*]. These types of studies could be extended to brain regions such as the lateral hypothalamus, which provides orexin/hypocretin inputs to the ventral tegmental area, which in turn project to the NAc. Animal models are also particularly conducive to determining if restoration of normal sleep patterns—for example, by using approaches (e.g., medications or neural stimulation) to promote sleep during times when it is abnormally deficient—reduces anhedonia. Such studies may be best conducted within an experimental design enabling analysis of sleep during the period of PRT testing, which was not performed in the present studies because we prioritized examining the effects of ELS on sleep patterns during adulthood. Collectively, these future approaches are consistent with precision medicine strategies to target specific symptoms in the subjects who are experiencing them. Regardless, an improved understanding of the relationship between sleep and the symptoms of depressive illness may enable rapid and transformational improvements in mental health outcomes.

